# ANOTHER TOOL IN THE TOOLBOX: *Wolbachia*-mediated protection against a specialized fungal pathogen of aphids

**DOI:** 10.1101/2023.07.24.550390

**Authors:** C.H.V. Higashi, B. Kamalaker, V. Patel, R. Inaganti, A. Bressan, J.A. Russell, K.M. Oliver

## Abstract

Aphids harbor nine heritable facultative symbionts, most mediating one or more ecological interactions. However, one aphid symbiont, *Wolbachia*, has eluded functional characterization despite being well-studied in other arthropods. In *Pentalonia* aphids, global pests of banana, *Wolbachia* was hypothesized to function as a co-obligate symbiont alongside the traditional obligate *Buchnera*, but subsequent genomic analyses failed to support this role. Sampling across multiple aphid populations, we found that > 80% of *Pentalonia* aphids carried a M-supergroup strain of *Wolbachia* named *wPni*. While the lack of fixation confirms that *Wolbachia* is not a co-obligate symbiont, the high infection frequencies in these entirely asexual aphids strongly suggested *Wolbachia* confers net fitness benefits. Finding no correlation between *Wolbachia* and aphid food plants use, we challenged *Wolbachia*-infected aphids with common natural enemies. While *Wolbachia* did not protect aphids against parasitoids, this symbiont conferred significant protection against the specialized fungal pathogen, *Pandora neoaphidis,* and improved aphid fitness in the absence of enemy challenge. Thus, we identified a new phenotype for the multifaceted *Wolbachia* and highlight a system that provides unique opportunities to merge key models of heritable symbiosis to better understand infection dynamics in nature and mechanisms underpinning symbiont-mediated phenotypes.

IMPORTANCE: *Wolbachia* symbionts spread and persist in a wide range of arthropods and nematodes by using a range of functional strategies, including reproductive manipulation, providing protection against viral and bacterial pathogens or by provisioning nutrients. Despite being one of the best-studied symbionts, little is known about the strains that infect aphids. In this study, we characterized the functional role of a Supergroup M *Wolbachia* strain infecting strictly asexual aphids of the genus *Pentalonia*. We report for the first time that this symbiont also defends against fungal entomopathogens and expand on the range of phenotypes conferred by this multifaceted symbiont.

## Introduction

Insects and other invertebrate animals frequently engage in heritable symbiosis with microbes (Moran et al. 2008; Douglas 2015). Heritable symbionts are primarily transmitted maternally and spread within host populations by providing net fitness benefits or manipulating host reproduction to increase the proportion of infected matrilines at the expense of uninfected ones (Werren et al. 2008, Oliver et al. 2014; Zug and Hammerstein 2015, Pietri et al. 2016). The most prevalent, taxonomically widespread, and versatile heritable symbiont on the planet is *Wolbachia* (Rickettsiales; α-Proteobacteria) (Zindel et al. 2011, Zug and Hammerstein 2012, Weinert et al. 2015, Landmann 2019). As a facultative symbiont, *Wolbachia* has been given the moniker ‘master manipulator’ as the only symbiont known to induce all four reproductive manipulation: parthenogenesis induction, cytoplasmic incompatibility, male killing and feminization (Werren et al. 2008; Kaur et al. 2021). Facultative *Wolbachia* are also associated with providing hosts with resistance to a wide range of pathogens, including viruses, bacteria, eukaryotic microbes, even nematodes (Saridaki and Bourtzis 2010, Zug and Hammerstein 2015, Correa et al. 2016, Sanda 2016). For instance, *Wolbachia* infecting *Drosophila* provides protection against a range of pathogens in both native and novel dipteran hosts (Hamilton and Perlman 2013, Pimentel et al. 2021; Kaur et al. 2021). In some cases, *Wolbachia* has become essential for host development (Dedeine et al. 2001; Comandatore et al. 2015), including as the B vitamin-provisioning obligate symbiont of blood-feeding *Cimex* bedbugs (Hosokawa et al. 2010; Nikoh et al 2015). It is further hypothesized that facultative *Wolbachia* strains may have more widespread roles as nutritional mutualists (Newton and Rice 2020).

*Wolbachia* are genetically diverse and grouped into at least 17 supergroups (A-F and H-R). Some supergroups are poorly characterized, indicating that novel roles for this symbiont are likely to be discovered. For example, while aphids (Hemiptera: Aphididae) have been important models for the study of symbiont-conferred phenotypes (Oliver et al. 2014), little is known about the role of *Wolbachia* in aphids, including strains from the M supergroup which appear restricted to aphids (Moreira et al. 2019; Romanov et al. 2020). This is surprising given that *Wolbachia* is relatively common in aphids (Augustinos et al. 2011, Wang et al. 2014, Gauthier et al. 2015, Chen et al. 2019, Guo et al. 2019, Ren et al. 2020; Romanov et al. 2020).

*Wolbachia* has been found in the banana aphid *P. nigronerversa* and its sole congener *P. caladii* (Jones et al. 2011; De Clerck et al. 2014), which vector *Banana bunchy top virus* (Nanoviridae); a major disease of banana and other musaceous plants (Hu et al. 1996, Kumar et al. 2015, Qazi 2016). These aphids are entirely asexual, indicating *Wolbachia* persistence likely results from mutualism rather than reproductive parasitism. One study hypothesized *Wolbachia* has become a nutrient-provisioning co-obligate symbiont, supporting decayed *Buchnera* function in *P. nigronervosa* (De Clerck et al. 2015). However, a reanalysis found little support for this co-obligate hypothesis (Manzano-Marin 2020), leaving open the role of *Wolbachia* in *Pentalonia* aphids.

In this study, we used a multi-pronged approach to better understand the role of *Wolbachia* in *Pentalonia* aphids. We first used 16S rRNA amplicon sequencing to identify the symbionts associated with *Pentalonia*, and diagnostic PCR to estimate infection frequencies across diverse populations. We found that *Wolbachia* was the only common aphid facultative symbiont, with high (but not fixed) infection frequencies confirming that *Wolbachia* is not a co-obligate nutritional symbiont in this system. Given that other aphid symbionts confer protection against fungal pathogens and parasitoids (Guo et al., 2017), we assayed *Wolbachia* protection against these threats. *Wolbachia* did not protect aphids against the parasitoid *Aphidius colemani*, nor a generalist fungal pathogen *Beauvaria bassiana*, but this symbiont did protect banana aphids challenged with the entomopathogenic fungus *Pandora neoaphidis*. This represents a new phenotype for *Wolbachia* that is potentially important across diverse arthropod host species.

## Methods

### Assessment of symbiont diversity in *Pentalonia* aphids

A total of 289 field colonies of *Pentalonia* aphids were sampled across the main Hawaiian Islands and Gainesville, Florida between 2011-2017 (Table *S1*) from four food plants (banana, heliconia, taro, ginger). Wingless individuals within the same colony cluster were sampled to maximize the likelihood of collecting clonally produced descendants of the same genotype. Aphids were placed into 1.5 ml tubes containing 95% ethanol and stored at −20°C until further use. Additional samples of *P. nigronervosa* collected on banana from Australia, Guam, and India was also obtained through collaborative efforts (see acknowledgments).

Given the morphological similarity of the two *Pentalonia* species, we amplified a diagnostic portion of the Cytochrome Oxidase subunit I (COI) gene to confirm species identity. Total DNA was extracted from individual, field-collected, adult aphids using a slightly modified Cetyltrimethylammonium bromide (CTAB) method (Doyle and Doyle 1990). Aphids were macerated in 200µl of CTAB buffer and incubated at 65°C for ≥ 1hr, then an equal volume of chloroform was mixed in and samples were centrifuged at 16,000g for 15 minutes. The supernatant was transferred to new 1.5ml centrifuge tubes and an equal volume of cold isopropanol was mixed in gently by inverting the tubes several times. After incubation at −20°C overnight, samples were thawed, then centrifuged for 20 min at 16,000g to pellet DNA. DNA was washed twice with 70% EtOH then a third time with 95% EtOH and allowed to air dry for ∼4hrs before rehydration in low TE buffer. This DNA template was used in PCR reactions to amplify a portion of the COI gene using primers LepF1 and LepR1 following the protocol described in Foottit et al. (2010). We then Sanger sequenced this amplicon in both directions and distinguished the two *Pentalonia* species by comparing sequences of our field-collected aphids to reference sequences (Foottit et al. 2010) using a Kimura 2 parameter model and visualized with a neighbor-joining tree.

To identify the full suite of endosymbionts present in *Pentalonia* aphids, we conducted 16S rRNA amplicon sequencing. Total DNA was extracted using the DNeasy Blood and Tissue kit (Qiagen) following the manufacturer’s protocols from pools of 30 randomly chosen, field-collected aphids of each *Pentalonia* species from five Hawaiian Islands, as well as *P. nigronervosa* from Australia and Guam. Extracted DNA was used as a template in PCR to target/amplify the hypervariable V3f (5’ CCTACGGGAGGCAGCAG 3’) and V4r (5’ GGACTACHVGGGTWTCTAAT 3’) regions of the bacterial 16S rRNA gene. Initial PCR amplifications were carried out in a 30 μl mixture that included 5 μl of KAPA HiFi Fidelity Buffer (New England Biolabs, UK), 0.3 μM of forward and reverse primers, 0.3mM each dNTPs, 0.5U of KAPA HiFi HotStart DNA Polymerase, 10 ng of template DNA and nuclease-free water up to 30 μl. The PCR conditions were 98 °C for 1 min (1 cycle), 98 °C for 20s, 55°C for 15 s and 72 °C for 45 s (30 cycles), followed by 72 °C for 7 min. The amplified products were checked by electrophoresis in 1% agarose gel, purified using the Cycle Pure Kit (Omega Bio-Tek), then quantified using a NanoDrop (Thermo Scientific) spectrophotometer. We used dual-index, paired read approach to sequence amplicons using the MiSeq 2 × 250 platform (Illumina, Inc. USA) at the Georgia Genomics and Bioinformatic Core (UGA, Athens, GA) in accordance with a previously described protocol (Kozich et al., 2013, Caporaso et al. 2012). Paired-end raw sequences were demultiplexed and merged based on overlaps using PEAR (Version 1.2.7) (Zhang at al., 2014). Reads were quality filtered using a quality filtering parameter (q value) Q20 and a p-value ≥90 to obtain high-quality reads using FastX Toolkit. Operational taxonomic units (OTUs) were constructed into clusters of greater than 97% sequence similarity using QIIME (Caporaso et al., 2010). Chimeras were detected based on the representative sequences of each OTU using UCHIME, and chimeric OTUs were excluded from the analysis (Edgar et al., 2011). Taxonomy was assigned to each OTU based on the Greengene database v13.5 reference database (DeSantis et al., 2006). Blank (i.e. template-free) controls were also included to detect contamination in laboratory reagents and extraction kits (Salter et al. 2014). OTUs present in these controls were excluded from analysis of aphid samples.

### Estimating *Wolbachia* infection frequency and characterizing strain diversity

Diagnostic PCR was used to estimate infection frequencies of *Wolbachia* across sampled populations. Using the CTAB method described above, total DNA was extracted from either single adult aphids (e.g. Florida samples) or pools of two clonal aphids (e.g. Hawaiian samples). We used three sets of *Wolbachia*-specific PCR primers to amplify fragments of three genes: 16S rRNA primers, W-specF and W-specR (Werren and Windsor 2000); glutamyltRNA amidotransferase (*gatB*) primers, gatB_F1/gatB_R1, and fructose-bisphosphate aldolase (*fbpA*) primers, fbpA_F1/fbpA_R1) (Baldo et al. 2006) (http://pubmlst.org/organisms/wolbachia-spp). Amplification conditions for the 16s rRNA primers included an initial pre-denaturation at 95°C for 30 s, followed by 30 cycles of 95°C for 30 s, 52°C for 30 s and 72 °C for 2 min, and ending with a final extension for 5 min at 72°C, while those for *gat*B and *fbp*A followed published protocols (Baldo et. al. 2006; http://pubmlst.org/organisms/wolbachia-spp). Samples that failedto amplify COI, potentially indicating poor template quality, were not screened for *Wolbachia*.All PCRs were performed in a final reaction volume of 20 μl including 4 μl of 5X reaction buffer (Promega, Madison, WI, USA), 1.5 μl of dNTPs (2.5 mM), 1.5 μl of forward and reverse primers (10 pmol), 1.2 μl of MgCl_2_ (25 mM) and 0.1 μl of Taq polymerase (Promega, Madison, WI, USA, 1U/μl), 8.7 μl H_2_O, and 2 μl of DNA. PCR products were visualized on 1.2 % agarose gels.

We also characterized *Wolbachia* strain diversity by obtaining a near full-length sequence of the 16S rRNA gene, using *Wolbachia*-specific 16S primers (see above) combined pairwise with “universal” bacterial reverse and forward primers 1507R and 10F (Moran et al., 2003; Werren and Windsor 2000). We also sequenced the five *Wolbachia* MLST typing loci *gatB*, *coxA, hcpA*, *ftsZ* and *fbpA*. Primers and PCR conditions for MLST loci can be found in (Werren and Windsor 2000; Baldo et al. 2006;) and PubMLST (https://pubmlst.org/organisms/wolbachia-spp). The published *ftsZ* primer failed to amplify the target, so we designed a new reverse primer (ftsZ_FMR2 5’-TTCCYGYGCRCCTTTCATTG-3’) that amplifies a ∼500 bp fragment that covers 100% of the typing region using the same reaction conditions. PCR amplicons for all loci were Sanger sequenced in both directions. All sequences generated were deposited in the GenBank database under accession numbers KJ786944 - KJ786953. MLST sequences were compared with those existing in the PubMLST *Wolbachia* MLST database (http://pubmlst.org/organisms/wolbachia-spp/). New allele sequences were submitted directly to the database curator and were assigned a new allele number (Table 4).

### Estimates of Maternal transmission rates

Maternal transmission rates of *Wolbachia* were estimated for *P. nigronervosa* lines collected from Florida (PP3) and Hawaii (Koloa). These lines were continuously maintained on banana leaf disks under laboratory conditions (20℃ ±1℃; 16 hr day:8 hr night light cycle). Individual offspring (in groups of 10-20 4^th^ instar to adult aphids) produced from *Wolbachia*-infected mothers were screened for *Wolbachia* using the CTAB DNA extraction protocol and diagnostic PCR as described above. Aphids were sampled over a 6-month period, which is approximately 16 generations given time from birth to first reproduction is approximately 11 days and the average life span of *P. nigronervosa* is roughly 25 days (Higashi, Unpublished). The aphid COI gene (see above) was used as a positive control for template viability.

### Phylogenetic analysis

16S rRNA sequences representing known super groups of *Wolbachia* were obtained from GenBank (accession numbers in Table S3) and sequences for *coxA, hcpA, fbpA, ftsZ,* and *gatB* were downloaded from the *Wolbachia* MLST website http://pubmlst.org/organisms/wolbachia-spp/) for use in phylogenetic analyses. Alignments were conducted using MUSCLE (Edgar, 2004) then visually inspected and manually adjusted as needed in Geneious v.10 (http://www.geneious.com/).

Maximum-Likelihood analyses were performed on the 16S rRNA sequence dataset and the concatenated MLST gene using the Randomized Axelerated Maximum Likelihood (RAxML) software v7.2.6 (Stamatakis et al., 2006) performed on the T-REX web server (Boc et al., 2012) (http://www.trex.uqam.ca/index.php?action=inference&project=trex). For both datasets we selected the GTR CAT evolutionary model and used the default Hill-Climbing algorithm available in RAxML. To analyze the inference robustness at the nodes of the trees we performed rapid alternative runs on 200 distinct starting trees with 1,000 rapid bootstrap random seed.

Bayesian analyses was performed on both 16S rRNA sequence and MLST datasets using MrBayes v3.1.2 (Huelsenbeck and Ronquist, 2001). The optimal evolutionary model for each sequence dataset selected using the Akaike Information Criterion in jModel Test v0.1.1 (Posada, 2008) resulting TVM+I+G for 16S rRNA, TVM+G for the *cox*A, GTR+I+G for the *fbp*A and the *gat*B, TIM3+I+G for the *fts*Z, and GTR+G for the *hcp*A. Posterior probabilities were approximated by sampling the trees using a Markov Chain Monte Carlo method and model parameters were set as priors in MrBayes. We kept the default parameters provided by the software and ran six million generations for all the datasets. We discarded the first 25% of tree samples from the cold chain (burn in) when computing 50% majority-rule consensus trees.

### Creating experimental aphid lines to examine the effects of *Wolbachia* infection

To understand *Wolbachia’s* role in banana aphids, we first established experimental lines with and without *wPni*. Unlike in other aphid models (e.g. Higashi et al., 2020), we were unable to cure banana aphids of *wPni* despite extensive efforts. Earlier efforts to cure *wPni* were also unsuccessful (De Clerck et al.,2015). Instead, we isolated and established clonal lines from two populations (FL and HI) that varied in *Wolbachia* presence. Based on the biology of this aphid, we expected the genotypes of aphid lines established from each locale to be highly similar, if not clonal. To confirm genetic similarity, we conducted microsatellite genotyping among these eight lines using five loci and using conditions as described in Galambao 2011. PCR products were then sent to Georgia Genomics and Bioinformatic Core (UGA, Athens, GA) as well as University of Pennsylvania Genomic and Sequencing Core (UPenn, Philadelphia, PA) for genotyping analysis on Applied Biosystems 3730xl and 3130xl DNA analyzer respectively, using the ROX500 size standard, and then analyzed using Geneious v.11 As predicted, we found few differences among amplified loci (Table S5) indicating our experimental lines shared highly similar genotypes. We also used diagnostic PCR (see above) to confirm that *wPni*+ aphids contained only *Wolbachia* (i.e. no other facultative endosymbionts) and that *wPni*-aphidsharbored only *Buchnera* (i.e. no facultative symbionts). *Wolbachia* presence/absence was similarly confirmed prior to all bioassays. Colonies of experimental lines were maintained on whole banana plants in BugDorm nylon mesh cages (#4F2260) at 22 ± 1°C under natural light conditions.

### Does *Wolbachia* protect banana aphids from the parasitoid *Aphidius colemani*?

We first used our experimental lines to test whether *Wolbachia* protects banana aphids against parasitoids as seen for several other aphid symbionts (Oliver et al. 2003, Vorburger et al. 2010, Heyworth and Ferrari 2015, McLean et al. 2020). *Aphidius colemani* (Hymenoptera: Braconidae) is a solitary parasitoid that attacks numerous aphid species including *Pentalonia* (Stary 1975, Völkl et al. 1990, Ode et al. 2005). Newly enclosed *A. colemani*, purchased from Beneficial Insectary© (Aphidiusforce C^®^), were provided honey and water and maintained in a biological incubator under 20°C and long light hours (16 light: 8 dark). Eight groups of ten 3^rd^-4^th^ instar aphids (8 reps/line; 80 total/line) from each aphid line were placed onto separate banana leaf disks. Leaf disks were prepared by cutting a circular portion of banana leaf (∼20mm diameter) and embedding in 1% agar within a small petri dish (35 x 10mm). A single mated *A. colemani* female was allowed to parasitize each aphid once and was removed from the leaf disk once all aphids were attacked. Parasitized aphids were maintained at 24 ± 1°C and the number of aphids that survived, mummified or had died (considered here as dual mortality because both aphid and wasp perished) were recorded 12 days post-parasitism. A generalized linear model (GLM) with binomial distribution and logit link function was used to compare parasitism outcomes across experimental lines (JMP Pro v.14 SAS Institute Inc., Cary NC). Pairwise contrasts between aphid lines were made using the contrast function after GLM.

### Does *Wolbachia* protect *Pentalonia* against fungal pathogens?

Given that multiple aphid symbionts confer protection against specialized fungal pathogens (Łukasik et al. 2013b, Parker et al. 2013), we first tested whether *wPni* protects against the aphid specialist fungal pathogen *P. neoaphidis*. To do this, previously preserved aphid cadavers infected by *P. neoaphidis* (USDA stain #2588) were rehydrated on 1.5% agar plates (35mm x 10mm) in the dark at 20°C for at least 14 hours to initiate sporulation. Petri dishes containing 2 *P. neoaphidis* sporulating cadavers were then inverted above a 35-mm petri dish containing 80 apterous 3rd-4th instar aphids from each of the eight *P. nigronervosa* lines. Aphids were exposed to *P. neoaphidis* for 2 hours and fungal plates were rotated between replicates of the same treatment every ∼30 minutes to normalize sporulation exposure of each aphid. Additionally, eight replicates (n=80) of the *P. neoaphidis* susceptible *Acyrthosiphon pisum* (pea aphid) line, WI-48, were included as a control to ensure our enemy challenge assay had worked. After exposure, banana aphids were transferred in groups of ten (8 reps/line) onto healthy banana leaf disks (as previously described) and pea aphid cohorts were transferred onto fresh *Vicia faba* (fava bean) plants. Both leaf disks and plants were placed in Percival incubators at 20°C at 100% humidity for 24hrs to induce sporulation.

We next determined whether *wPni* protects against the entomopathogenic generalist, *Beauvaria bassiana,* a commercially available bioinsecticide used to control a variety of insect pests. *Beauvaria* strain GHA (BotaniGard® 22WP) was mixed into solution using distilled water following the manufacturer’s small volume rate. Individual 3rd-4th instar aphids were dipped and swirled in the *Beauvaria* solution serval time for approximately 3-5 seconds and then immediately transferred onto a healthy banana leaf disk. A total of ten *Beauvaria* exposed aphids were placed on a single health banana leaf disk (= 1 rep) and a total of 5 replicates were established for each of our eight *P. nigronervosa* lines. Aphids were monitored every 24 hours for 10 days to assess aphid survival, mortality and fungal sporulation. Infected corpses were removed to prevent secondary fungal infections. We removed offspring periodically to minimize crowding and changed leaf disks on an as-needed basis. A generalized linear model (GLM) with binomial distribution and logit link function (JMP Pro v.14, SAS Institute Inc., Cary NC) was used to compare survival, mortality and sporulation proportions within (*wPni*- or *wPni*+) and between (*wPni*-vs *wPni*+) aphid groups. Pairwise contrasts between aphid lines were made using the contrast function after GLM.

### Fitness effects associated with *Wolbachia* in the absence of enemy challenge

To assess constitutive fitness cost to harboring *Wolbachia*, we measured aphid development time (birth to adulthood), cumulative fecundity and longevity (measured as 50% survival) for our *wPni*- and *wPni*+ aphid lines. Fifty to sixty adult aphids of each line were placed onto large banana leaf disks (100mm x 15mm) and allowed to reproduce for 24 hrs. Groups of 10 first instar aphids from each line were then placed onto small banana leaf disks (= 1 replication). Leaf disks and aphids were maintained at 24℃ ± 1℃ and checked for molting every 2 days (an indication the aphid has transitioned to the next developmental stage) until adulthood. Aphids were transferred to fresh banana leaf disks approximately every 4-5 days as needed. A nonparametric Wilcoxon rank sum test used to compare developmental times among lines with the same infection status and between *wPni*- and *wPni*+ aphids.

Cumulative fecundity and 50% survivorship were assessed by allowing five newly developed adults from each *wPni*- and *wPni*+ lines to reproduce (=1 rep) on fresh banana leaf disks at 25℃ +/−1. The number of offspring produced was counted every 2-3 days and then removed. Adult aphids were transferred to new banana leaf disks approximately every 4-5 days as needed. Aphid mortality was recorded and offspring counts continued until all the adults had died. Cumulative fecundity was compared within groups using analysis of variance (ANOVA) with Tukey’s honestly significant difference (HSD) test. Fecundity between *wPni*- and *wPni*+ aphids was compared using a Student’s 2-tailed t-test. Aphid survivorship was analyzed using a nonparametric Wilcoxon rank sum test. Aphid survival was fit to a Weibull distribution to estimate α=50% survival time.

## Results

**Hawaiian *Pentalonia* aphid populations have low symbiont diversity but are frequently infected with *Wolbachia*.** We detected four heritable endosymbionts inhabiting *Pentalonia* across the seven pooled samples analyzed in our 16S amplicon sequencing (Figure 1). As expected, the obligate symbiont *Buchnera* was found in all *Pentalonia* samples and often exhibited the greatest read abundance (range 10.13% - 91%). *Wolbachia* was the only facultative symbiont detected in both *Pentalonia* species from all sampled locations (Figure 1). The facultative symbiont *Serratia symbiotica* was the only other symbiont detected in both *Pentalonia* species, but found on only three of the five islands sampled. Sequencing and conducting BLASTn analysis (Altschul et al. 1990) of portions of the *accD*, *gyrB*, and *recJ* genes showed that the *S. symbiotica* isolate from *Pentalonia* was 99% similar to strains found in other aphids (GenBank accession # OR283239 - OR283241). We detected *Arsenophonus* only in *P. nigronervosa* collected from the island of Moloka’i. Sequencing a 500 bp portion of the 23S ribosomal RNA gene revealed that *Arsenophonus* from *Pentalonia* (accession #OR271588) was most similar to strains isolated from honeybees (∼99%) and whiteflies (∼97%) (see *Table S2* for all loci and primers).

**Figure 1:**
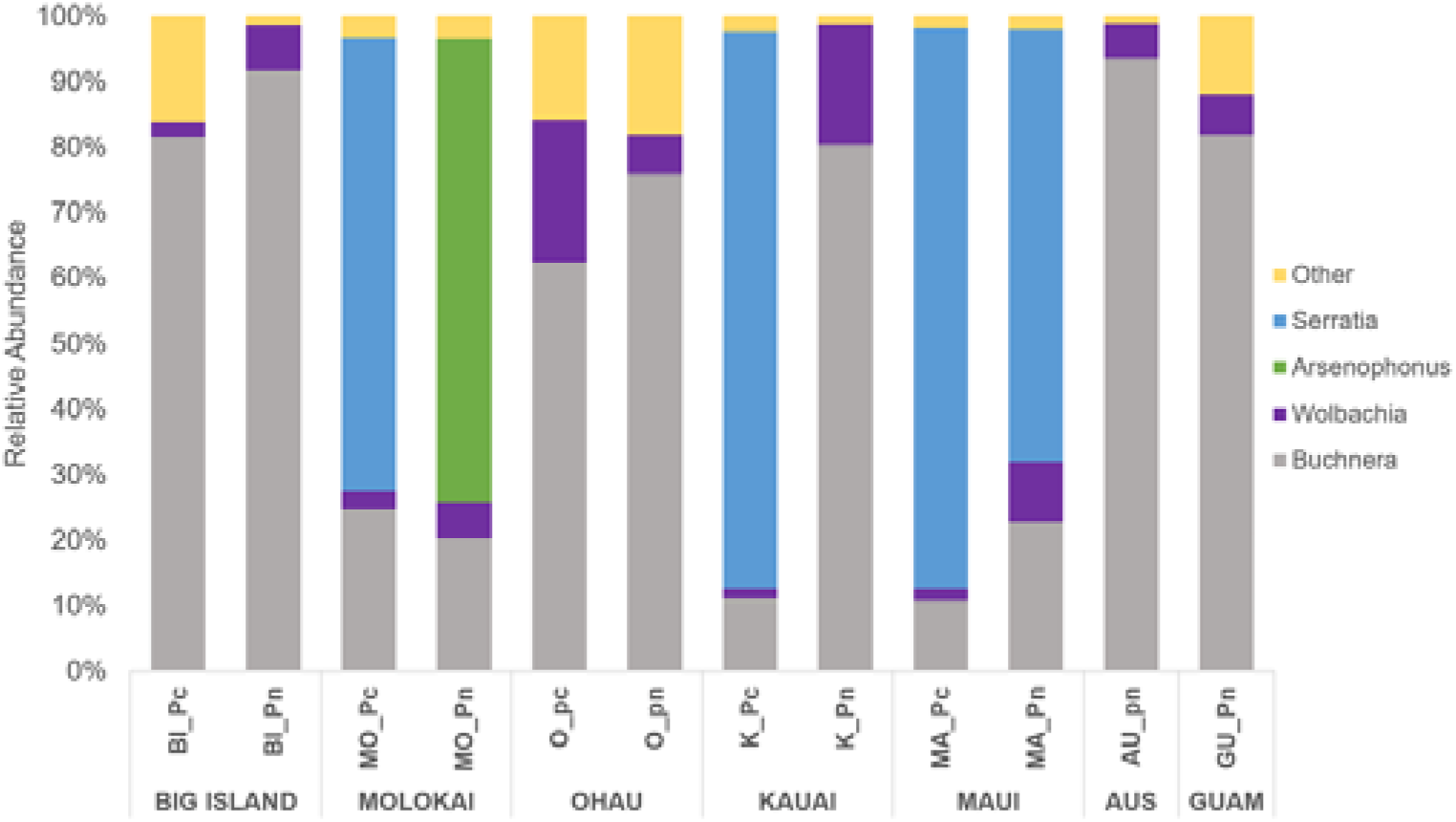
Relative read abundance (%) of endosymbiotic bacteria detected in pooled samples of *P. nigronervosa* (Pn) and *P. caladii* (Pc) collected across five Hawaiian Islands, Australia (AUS) and Guam (GU) using 16s rRNA. *Other = lineages predicted to be environmental associates, not heritable symbionts.

***Wolbachia* infection is high but not fixed in natural populations.** DNA was successfully extracted from 261 out of 282 *P. nigronervosa* and *P. caladii* sampled clones across five Hawaiian Islands. Overall, *Wolbachia* occurred in 83% (217/261) of *Pentalonia* sampled. Rates of infection did not differ significantly between *P. nigronervosa* and *P. caladii* (83.3% and 83.0%, respectfully; Table 2). Infection rates among *P. nigronervosa* across the five Hawaiian Islands were also statistically similar, however, infection rates varied significantly among the island populations for *P. caladii;* Molokai had the lowest rate of infection at 59% and Big Island had the highest infection rate at 90% (2-tailed fisher’s exact test*; P* < 0.001; Table 2).

**Table 2:** Incidence of *wPni* infection across sampled *Pentalonia* colonies.

| Location | Colonies Screened | Positive for *Wolbachia* | Avg. Infection Rate |
| --- | --- | --- | --- |
| *P. nigronervosa* * |  |  |  |
| Kauai | 23 | 19 | 82.6% |
| Oahu | 7 | 6 | 85.7% |
| Maui | 39 | 31 | 79.5% |
| Molokai | 17 | 13 | 76.5% |
| Big Island | 34 | 31 | 91.2% |
| **Total** | **120** | **100** | **83.3%** |
| *P. caladii*† |  |  |  |
| Kauai | 26 | 22 | 84.6% |
| Oahu | 16 | 12 | 75.0% |
| Maui | 44 | 39 | 88.6% |
| Molokai | 17 | 10 | 58.8% |
| Big Island | 38 | 34 | 89.5% |
| **Total** | **141** | **117** | **83.0%** |
| *Overall* * |  |  |  |
| Kauai | 49 | 41 | 83.7% |
| Oahu | 23 | 18 | 78.3% |
| Maui | 83 | 70 | 84.3% |
| Molokai | 32 | 23 | 71.9% |
| Big Island | 72 | 65 | 90.3% |
| **Hawaii Total** | **261** | **217** | **83.1%** |

\* Not significantly different by 2-tailed fisher's exact test.
† Infection rates determined significantly different by 2-tailed fisher's exact test.

**Table 3:** Incidence of *wPni* infection in *Pentalonia* aphids found on various food plants.

| Plant Family | Pn<br>(# of W+ /<br># screened) | Pc<br>(# of W+ /<br># screened) | Pn+Pc<br>(# of W+ / #<br>screened<sup>*</sup>) | % |
| --- | --- | --- | --- | --- |
| Musaceae<sup>a</sup> | 90 / 110 | 1 / 1 | 91 / 111 | 82.0 |
| Araceae<sup>b</sup> | 7 / 7 | 11 / 13 | 18 / 20 | 88.9 |
| Zingiberaceae<sup>b</sup> | 2 / 2 | 92 / 111 | 94 / 113 | 83.2 |
| Heliconiaceae<sup>b</sup> | 1 / 1 | 8 / 10 | 9 / 11 | 81.1 |
| Costaceae<sup>b</sup> | 0 / 0 | 5 / 6 | 5 / 6 | 83.3 |
| % | 83.3 | 83.0 | 83.1 |  |
| P-value<br>(Chi-sq) | 0.54<br>(2.18) | 0.99<br>(0.29) | 0.94<br>(0.79) |  |

*PN* = *P. nigronervosa*; *PC* = *P. caladii*
<sup>\*</sup> = DNA extracted from 2 aphids sampled from a single colony on a single host plant.
<sup>a</sup> = Banana plants mainly colonized by *P. nigronervosa*
<sup>b</sup> = Plant hosts mainly colonized by *P. caladii*

*Pentalonia caladii* and *P. nigronervosa* have distinct food plant preferences. *P. nigronervosa* feeds primarily on plants in the Musaceae, including banana, while *P. caladii* feeds on plants from several families, including the Araceae (Taro, Elephant ear), Zingiberaceae (Ginger), Heliconiaceae (Heloconia), and Costaceae (Spiral Ginger) (Foottit et al. 2010; Rahmah et al. 2021). This food plant association was observed in our field collections (Fisher’s exact test; *P* < 0.001; Table S6). However, we did not find significant associations between *Wolbachia* prevalence and food plant use within or between *Pentalonia* species (2-tailed fisher’s exact test; *P* > 0.05, Table 3).

A single, novel, M supergroup *Wolbachia* strain was found across most *Pentalonia* populations and is vertically transmitted at high rates. A single *wPni* strain, identical at all five loci, was found in all individuals tested (n=214) from both *Pentalonia* species sampled across the five Hawaiian Islands. This novel strain was submitted to the MLST database and assigned a new strain type (ST#402; Table S4). Sequences for *gatB*, *coxA*, *hcpA*, and *ftsZ* genes were assigned new allele numbers except for *fbpA* which matched an existing allele #344 (Table S4). The single strain (ST#402) recovered from the Hawaiian populations was also found in *P. nigronervosa* from Australia and Florida. *wPni* isolates from India and Guam shared allelic identity at four of the five loci sampled for ST#402 (Table S4) but were designated as different strain types, ST#405 and ST#406, respectively (Table 4).

Maximum likelihood phylogenies based on ∼1400bp of the16S rRNA gene showed *wPni* clustered within the M supergroup (Figure 2), a clade that, to date, is associated only with*Wolbachia* from aphids (Augustinos et al., 2011; Wang et al., 2014). An unrooted maximum likelihood phylogeny constructed from concatenated MLST sequences show all three *wPni* strains clustering together in support of the M supergroup placement (Figure S1).

**Figure 2:**
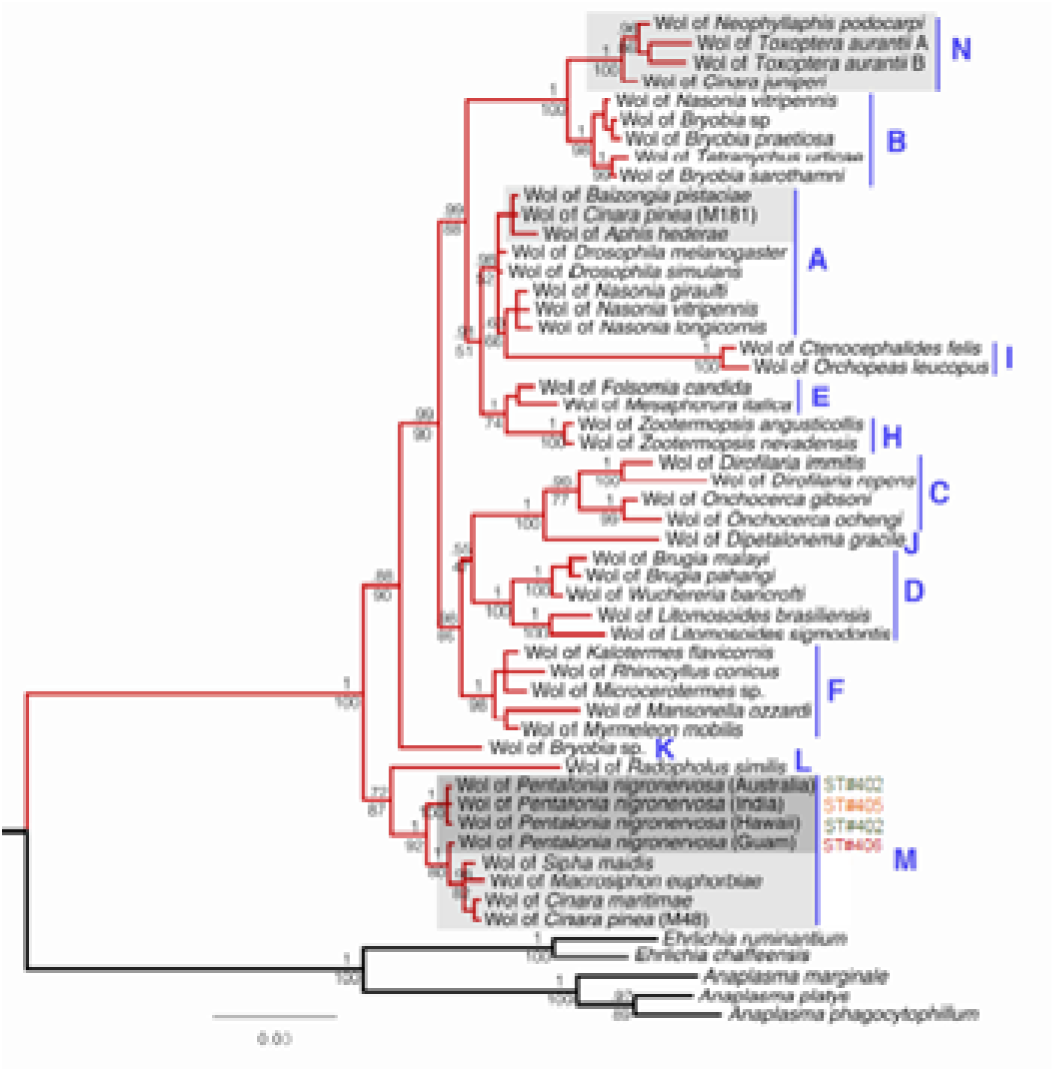
Phylogeny of *Wolbachia* (red branches) based on 1400bp of 16S rRNA with an emphasis on strains colonizing aphids of the genus *Pentalonia* (dark grey box) and other aphid species identified by Augustinos et al., 2011 (light grey boxes; Panel A). The 16S rRNA sequences of two Rickettsiales bacteria, *Anaplasma* and *Ehlirichia* (black branches) were used as outgroups. Capital blue letters and the vertical blue bars indicate *Wolbachia* super-groups.

The common *Wolbachia* ST#402 was also maternally transmitted at high rates in our laboratory assay. Transmission rates of *wPni* carried by two aphid lines, one from Florida (PP3; N = 102) and one from Hawaii (Wailua; N = 104) were 100% (Table S7) across approximately 16 generations.

Values above nodes are for the posterior probabilities generated from the Bayesian analyses. Values below the nodes are bootstrap values from the Maximum Likelihood analysis (ML for the tree: −6297.33). For accession numbers refer to Table S3. The strain type associated with *Pentalonia* populations are indicated to the right.

*Wolbachia* provides protection against the specialized fungal pathogen *Pandora neoaphidis*, but not against the generalist *Beauveria bassiana.* We observed low rates of aphid survival (10%) and high rates of fungal sporulation (46%) from our susceptible pea aphid control line (WI-48) when exposed to the aphid specialist pathogen *P. neoaphidis* (Figure 3A). This demonstrates that our enemy challenge assay exhibited pathogenicity levels similar to those of past experiments (Doremus & Oliver, 2017). When challenging banana aphids, we found that those with *Wolbachia* exhibited significantly higher rates of survival and lower rates of fungal sporulation compared to those without *wPni* (Figure 3A). Specifically, survival was 24% higher for *Pentalonia* carrying *wPni* compared to those lacking *Wolbachia* (*χ*^2^ *=* 45.43, P < 0.0001; Figure 3A), while the rate of fungal sporulation was 27% lower for those carrying the symbiont (*χ*^2^ *=* 51.80, P < 0.0001). No variation in aphid survival and sporulation was observed (GzLM, *df* = 3, P > 0.05; Figure 3B) among the four *wPni*-aphid lines, suggesting endogenous defenses did not vary among symbiont-free lines. In contrast, survival (GzLM; *χ*^2^ = 9.65, *df* = 3, P = 0.022) and fungal sporulation varied significantly (GzLM; *χ*^2^ = 36.61, *df* = 3, P < 0.0001) among the four *wPni*+ lines (Figure 3C).

**Figure 3:**
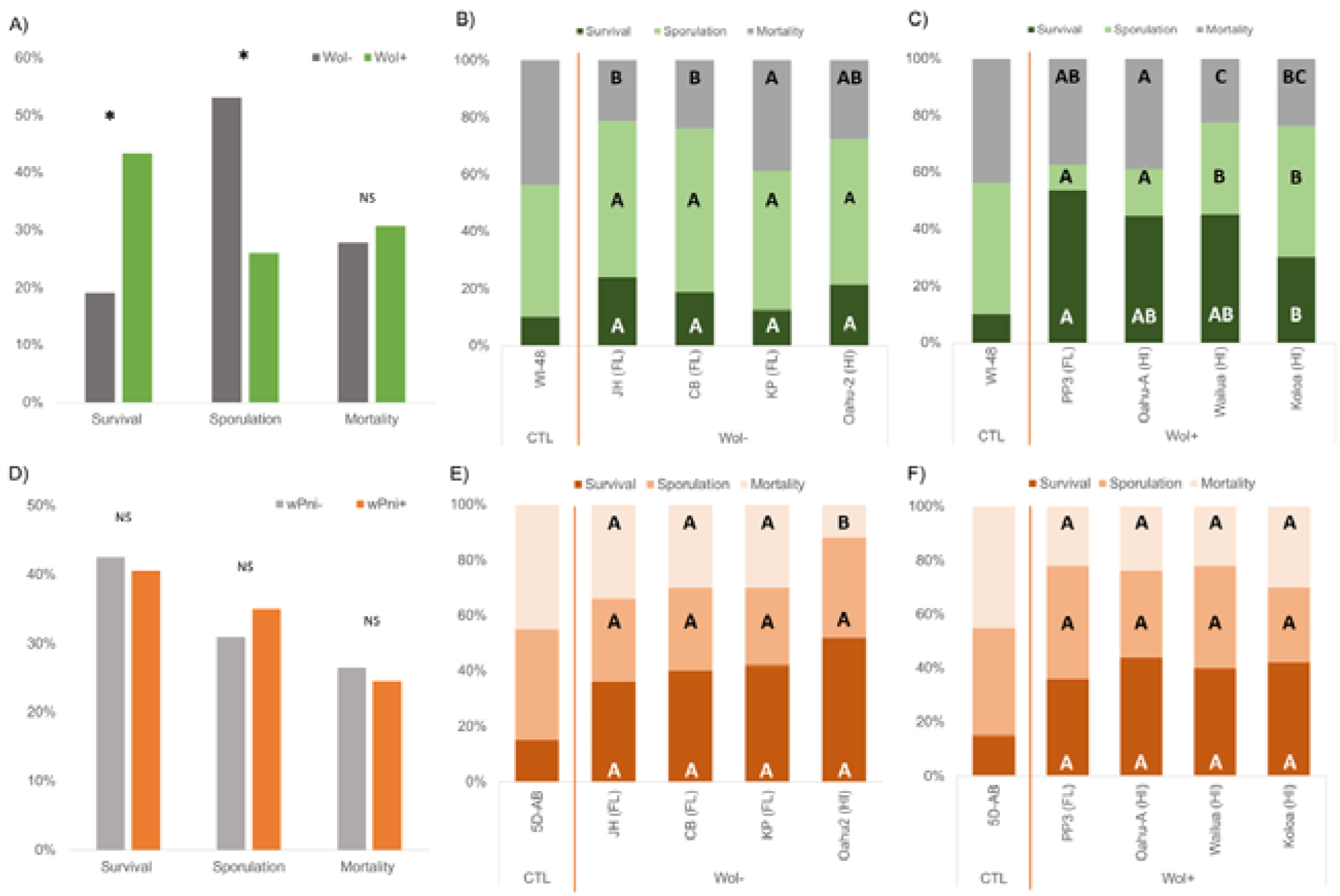
Aphid survival, fungal sporulation, and dual-mortality rates of *P. nigronervosa* aphid lines with (*wPni+*) and without (*wPni*-) 10-days post exposure to the entomopathogen (A-C) *P. neoaphidis* or (D-F) *B. bassiana*. The asterisk above bars in panels A and C indicate significant difference (P ≤ 0.05) and ‘NS’ indicates no significant difference (P > 0.05). Letters in panels B, C, E, & F indicates significant differences for each of the exposure outcomes among *wPni*-(B, E) and *wPni+* (C, F) aphids by pairwise contrasts after GzLM (NS = *P* > 0.05). Results from pea aphid control lines WI-48 (for *P. neoaphidis*) and 5DAB (*B. bassiana*) were only used as a visual reference and not included in the analysis of (B, E) or (C, F).

When exposed to the generalist pathogen *B. bassiana*, aphids with and without *wPni* experienced similar survival (GzLM; *χ*^2^ = 0.165, *df* = 1, P = 0.685) and fungal sporulation rates (GzLM; *χ*^2^ = 0.724, *df* = 1, P = 0.395) (Figure 3D). We also found no significant variation in aphid survival or fungal sporulation rates among banana aphids with *Wolbachia* or those lacking this symbiont (GzLM, *df* = 3, P > 0.05) (Figure 3E, F).

***Wolbachia* failed to protect banana aphids against the parasitoid *A. colemani.*** In our parasitoid challenge assays, rates of mummification, a common proxy for successful parasitism (Oliver et al. 2012), did not vary between *wPni*+ and symbiont-free banana aphid lines (Figure 4; GzLM *P* > 0.05). Given this result, it is unlikely that the small (∼5%), but marginally significant (GzLM; *χ*^2^ *=* 3.88*, df* = 1, P = 0.05) increases in aphid survival found for *wPni*+ lines reflect direct anti-parasitoid defenses. Instead, these modest increases in survival more likely reflect indirect effects resulting from increased dual-mortality (both aphid and wasp perish) (*χ*^2^ *=* 8.67*, df* = 1, P = 0.003) in parasitized symbiont-free aphids (Figure 4A). We recovered significant variation in parasitism outcomes among symbiont-free aphid lines (Figure 4B) and among lines carrying *Wolbachia* (Figure 4C) but the basis of this variation is unclear.

**Figure 4:**
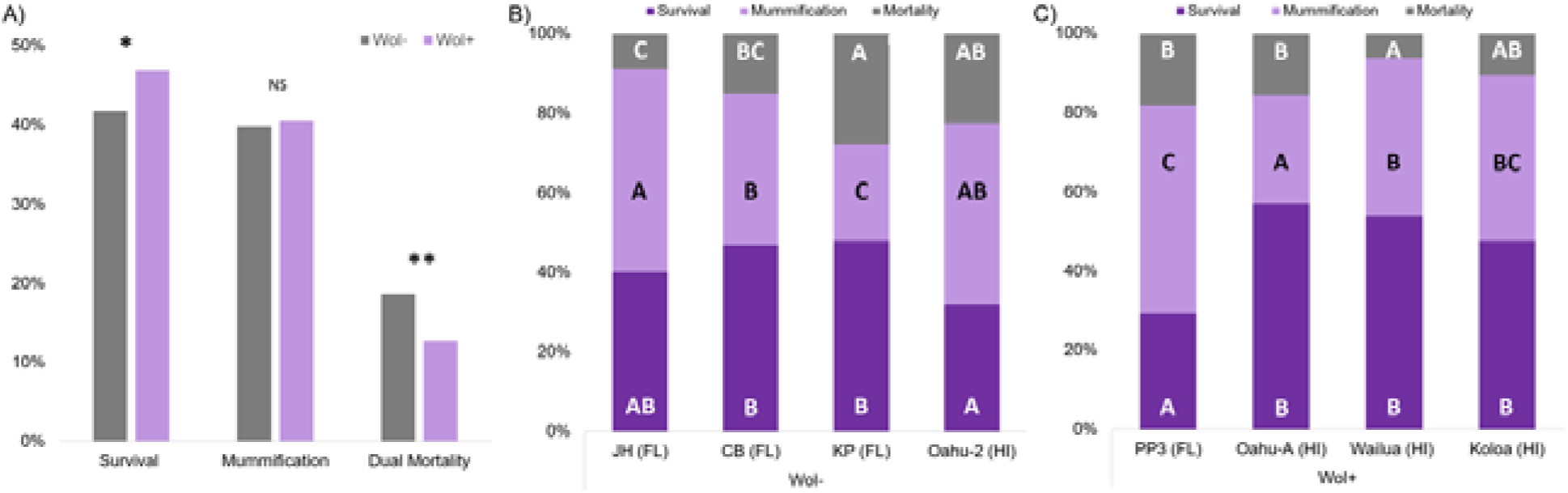
Outcomes 12 days after enemy challenge by the parasitoid *A. colemani*. (A) % of aphids surviving, % of aphids mummifying, and dual mortality (both aphid and wasp perish) between *wPni+* and *wPni-* lines. (NS = *P* > 0.05; * = *P* ≤ 0.05; ** = P ≤ 0.01). Also presented are within-group comparisons of outcomes among (B) *wPni*- and (C) *wPni+* aphids with letters indicating significant differences.

### *Wolbachia* confers net fitness benefits for banana aphids in the absence of enemy challenge

On average, *wPni+* aphids developed faster (*χ*^2^ = 12.8; P = 0.0003), produced more offspring (t = 1.74; P = 0.043) and lived significantly longer (*χ*^2^ = 16.3; P < 0.0001; Figure S2) than aphids without *wPni* (Table 5). Fitness performance varied little among the four *wPni*-aphid lines except for line ‘Oahu-W’ which showed a small but significant increase in survival relative to the other *wPni*-lines (Table 5). We observed differences in development time and fecundity between the four *wPni+* aphid lines (*χ*^2^ *=* 13.89, *df* = 3, P = 0.0031) (Table 5). The *wPni*+ aphid line ‘PP3 (FL)’ performed best in these metrics, while the ‘Koloa’ exhibited the lowest performance. Interestingly, these two lines represented the best and the worst protection against *P. neoaphidis,* respectively (Figure 3C).

**Table 5:** Mean (± SE) development time, cumulative fecundity and aphid longevity (50% survival) for each aphid line.

| Line (loc) | *wPni* + / - | Development Time hrs ± SE | Cumulative Fecundity Offspring ± SE | 50% Survival days |
| --- | --- | --- | --- | --- |
| Within Group |  |  |  |  |
| JH (FL) | *wPni* (-) | 278.3 ± 3.7 a | 53.5 ± 3.9 a | 19.7 a |
| CB (FL) | *wPni* (-) | 271.0 ± 3.3 a | 53.5 ± 11.4 a | 21.1 a |
| KP (FL) | *wPni* (-) | 273.0 ± 3.6 a | 60.1 ± 14.3 a | 24.2 ab |
| Oahu-W (HI) | *wPni* (-) | 277.6 ± 2.1 a | 56.4 ± 13.2 a | 25.1 b |
| Within Group |  |  |  |  |
| PP3 (FL) | *wPni* + | 260.1 ± 2.9 c | 71.9 ± 11.2 a | 24.2 a |
| Oahu-A (HI) | *wPni* + | 265.6 ± 2.8 bc | 66.0 ± 10.3 a | 27.4 a |
| Wailua (HI) | *wPni* + | 268.8 ± 3.2 ab | 60.3 ± 17.3 a | 27.3 a |
| Koloa (HI) | *wPni* + | 272.5 ± 2.7 a | 47.6 ± 1.8 b | 22.3 a |
| Between Groups |  |  |  |  |
| | *wPni* (-) | 275.1 ± 1.6 b | 55.88 ± 2.0 b | 21.8 b |
| | *wPni* + | 266.5 ± 1.5 a | 61.45 ± 15.8 a | 25.2 a |

<sup>†</sup> Development time was analyzed using multiple Wilcoxon Rank Sum tests.
<sup>‡</sup> Within-group fecundity was compared using a Tukey’s post hoc test. A one-tail t-test was used to compare *wPni*- and *wPni*+ groups ( $P > t$ ). Letter after value denotes significant difference at $P < 0.05$ .
<sup>§</sup> Survival data was fit to Weibull distribution for estimates of 50% survival. Letter after value indicates significant difference ( $P < 0.05$ ) by Wilcoxon test.

## Discussion

Aphids are leading models for characterizing symbiont-conferred phenotypes (Oliver et al 2010; Brisson and Stern 2006, Huang and Qiao 2014; Brandt et al 2017). Conspicuously absent, however, have been studies elucidating the roles of the phenomenally widespread and phenotypically diverse symbiont *Wolbachia* in aphids. While originally thought to be uncommon in aphids (Jeyaprakash and Hoy 2000), *Wolbachia* now appears to be a frequent passenger of this economically important insect group, including the presence of strains from supergroups M and N that appear restricted to aphids (Augustinos et al. 2011, De Clerck et al., 2014; Zytynska and Weisser 2016, Chen et al. 2019, Moreira et al. 2019; Wang et al., 2014). Here we examined *Wolbachia* carried by entirely parthenogenetic *Pentalonia* aphids, which are important pests of banana.

Using 16S rRNA amplicon sequencing of more than 360 aphids pooled across 12 populations, we found that that only three (*Wolbachia*, *Serratia symbiotica* and *Arsenophonus*) of the nine facultative symbionts that commonly infect aphids (Guo et al. 2017) were present in one or both *Pentalonia* species (Figure 1). This low diversity heritable microbiome contrasts with, for instance, the seven facultative symbionts routinely found in most sampled pea aphid populations (Russell et al. 2013a). The asexuality of *Pentalonia* may reduce opportunities for acquiring symbionts (Moran and Dunbar 2006).

*Wolbachia* was the only facultative symbiont present in all populations (Figure 1). MLST and phylogenetic analysis identified a single, widespread *Wolbachia* strain (ST#402) from the M supergroup (Table S4) present in all Hawaiian *Pentalonia* samples, and in limited sampling from Australia and Florida. This is consistent with the widespread dispersal of *Pentalonia* aphids via the banana and ornamental plant trades (Halbert et al. 2015). Similar, but distinct, *wPni* strains were found in *Pentalonia* populations from Guam (ST# 406) and India (ST# 405) indicating some strain variability is present. Anholocyclic aphids, such as *Pentalonia,* are often represented by a few dominant clonal genotypes (Galambao 2011; Harrison & Mondor 2011), which in turn may limit strain diversity of their inherited symbionts (Dykstra et al. 2014). Fitness benefits and efficient maternal transfer (100% in our lab-based assays; Table S7) likely drive the spread of specific strains, but occasional horizontal transmission via shared food plants or parasitoids may also contribute (Caspi-Fluger et. al. 2012, Geher & Vorburger 2012). More work is needed to better understand strain variation in this system and how strains interact with a presumably limited set of aphid genotypes (Parker et al. 2021).

As noted above, *Wolbachia* was first hypothesized to be a nutrient-provisioning co-obligate symbiont supporting degraded *Buchnera* function in *P. nigronervosa* (De Clerck et al. 2015). This hypothesis was based on findings that all individuals tested carried *Wolbachia,* the inability to selectively cure aphids of *Wolbachia*, and a reconstruction of metabolic pathways from a metagenomic assembly suggesting that *Wolbachia* are needed to produce key nutrients, including riboflavin, missing from the banana aphid’s *Buchnera*, and which are typically present in *Buchnera* of other species. While genome degradation in *Buchnera* and other obligate symbionts has driven the transition of numerous facultative symbionts toward co-obligate status (Bennett & Moran 2015, Meseguer et al. 2017), Manzano-Marin (2020) found little support for degraded *Buchnera* in *Pentalonia* as most of the genes originally thought to be missing were found to be intact. Our own PCR-based assays and Sanger sequencing confirmed the presence of the putatively missing loci in the B2 (riboflavin) pathway (data not shown). A more conclusive nail in the co-obligate hypothesis coffin, is that our survey clearly indicates that *Wolbachia* is a facultative symbiont in banana aphids. Infection frequencies of *Wolbachia* in *P*. *nigronervosa* across the Hawaiian Islands and other populations were high (*x―* = 83%; N = 120; Table 2), but never fixed; a requirement for co-obligate status. We found similar infection frequencies (*x―* = 83%; N = 141; Table 2) in *P. caladii*, consistent with an earlier report for this species (Jones et al. 2011) and suggestive of similar roles for *Wolbachia* in these related aphids. It is possible that *Wolbachia* supplied nutrition is conditionally required, and may, for example, be particularly beneficial on specific food plants. However, our surveys showed no association between *Wolbachia* presence and food-plant usage (Table 3).

The high infection frequencies, combined with high rates of maternal transmission in these exclusively parthenogenetic aphids strongly suggested *Wolbachia* confers net fitness benefits. Given that the other common aphid symbionts confer protection against specialized natural enemies (Oliver et al. 2003; Schmid et al. 2012; Asplen et al. 2014; Scarborough et al. 2005, Łukasik et al. 2013b, Parker et al. 2013, Heyworth and Ferrari 2015, Parker et al. 2021), we investigated whether *wPni* protects *Pentalonia* aphids against fungal entomopathogens and a common parasitoid. When challenged with the specialist fungal pathogen *P. neoaphidis*, banana aphids carrying *wPni* experienced higher rates of survival and lower fungal sporulation compared to those without the symbiont. In contrast, aphids with and without *wPni* were equally susceptible to the generalist *B. bassiana,* highlighting the specificity of *wPni* anti-fungal defense (Figure 2). While diverse beneficial roles have been identified for *Wolbachia* in the past decade, which include protection against diverse pathogens (Hedges et al. 2008, Teixeira et al. 2008, Fenton et al. 2011, Hughes et al. 2011, Zélé et al. 2012), this is the first report, to our knowledge, of *Wolbachia* conferring protection against a fungal entomopathogen. Specialized fungal pathogens are a persistent problem for insects, and many groups, including aphids, lack adequate endogenous defenses (Gerardo et al. 2010, Wang and Wang 2017). Thus, *Wolbachia*-mediated anti-fungal defenses are potentially widespread across insects and their relatives.

While all *Wolbachia*-carrying *Pentalonia* aphid lines were more protected than *wPni*-lines, the PP3 line had higher survival and lower sporulation then all others (Figure 3C). Symbiont strain variation has been shown to confer variable defensive phenotypes in other aphid systems (Martinez et al. 2014b, Cayetano and Vorburger 2015, Parker et al. 2017, Mathé-Hubert et al. 2019, McLean et al. 2020) and similar stain varibility also characterizes *Wolbachia*-conferred phenotypes in other insect systems (Bordenstein and Werren 2007, Graham et al. 2012, Chrostek et al. 2013, Martinez et al. 2014, Hoffmann et al. 2015, Martinez et al. 2017). While we observed no variation in symbiont genotypes for the isolates used in our bioassays, the factors responsible for phenotypic differences in protective symbioses are often located on mobile elements (Oliver et al. 2009, Chevignon et al. 2018, Lindsey et al. 2018a). Hence our multi-locus typing protocols may not capture relevant *wPni* strain variation. Unfortunately, our inability to manipulate *wPni* infections in the banana aphid system ultimately limits inferences about the sources of variation. However, the lack of variability in *Wolbachia-*free aphid lineages following *Pandora* challenge, suggests that symbiont strain, or symbiont genotype x host genotype interactions are more likely to contribute this variation (Vorburger et al. 2009, Parker et al. 2017, Weldon et al. 2020). While experimental manipulation among shared genotypes is the gold standard for isolating effects of symbiont infection (Oliver et al. 2010), the consistent anti-*Pandora* protection observed across multiple, highly similar aphid genotypes provides clear support for *Wolbachia*-mediated protection against specialist fungal pathogens. Despite six aphid facultative symbionts now shown to confer protection against fungal pathogens little is known about the mechanisms underlying this protection (Brownlie and Johnson 2009). The frequent occurrence of this phenotype among diverse symbiont lineages suggests that ‘immune priming’or competition for resources shared between host and symbiont may be the most likely general mechanisms, although production of anti-fungal peptides remains a possibility (Gerardo et al. 2020).

In contrast, *Wolbachia* did not protect aphids against attack by the parasitoid, *A. colemani* (Figure 3A). Several parasitoid species have been evaluated for control of natural populations of banana aphids (Völkl et al. 1990, Wellings et al. 1994), and it remains possible that *Wolbachia* protects against other wasp species. For example, strains of other aphid symbionts, including *Hamiltonella* and *Spiroplasma,* have been shown to protect against some parasitoid species, but not others (Asplen et al. 2014, Cayetano and Vorburger 2015, McLean and Godfray 2015, Martinez et al. 2016, McLean et al. 2020).

*Wolbachia* also improved aphid fitness in the absence of enemy challenge. *Pentalonia* aphids infected with *wPni* were more fecund, lived longer, and develop into adults faster than those without *wPni* (Table 5; Figure 3). Thus, not only does *wPni* protect *Pentalonia* against a common fungal pathogen, but also generally improves host fitness under permissive conditions. Previous studies indicate that protective symbionts in other aphid systems exert variable effects on hosts absent natural enemies, some imposing strong constitutive costs while other are relatively benign (Russell and Moran 2006, Vorburger et al. 2013, Weldon et al. 2013b, Polin et al. 2014, Doremus et al. 2017, Weldon et al. 2020). For example, some *R. insecticola* strains that protect against *P. neoaphidis* are relatively benign in the absence of fungal challenge. Another facultative symbiont, *Rickettsia,* in whiteflies, can manipulate reproduction and provide protection against bacterial pathogens, but also improves fitness in the absence of enemy challenge; although the latter has been shown to vary with host genotype (Himler et al. 2011, Hendry et al. 2014, Cass et al. 2016).

## Conclusions

*Wolbachia* is the best-studied symbiont with an ever-increasing repertoire of functions that support its spread and maintenance across diverse arthropod populations. In this study, we expand on the tools utilized by *Wolbachia* showing for the first time that this symbiont also defends against fungal entomopathogens. This is also the first report to our knowledge of function for an M supergroup strain of *Wolbachia,* which commonly occur in aphids. The *Pentalonia* aphid model with *wPni* as the single, dominant facultative symbiont, provides opportunities to merge two important models of symbiosis (aphids and *Wolbachia*) to better understand the biology of each. This system also provides unique opportunities to expand symbiont-assisted control of agents that threaten human interests. *Pentalonia* aphids are economically important vectors of *Banana bunchy top virus* (BBTV), the causative agent of Banana bunchy top disease which is a serious threat impacting banana production. Given that *Wolbachia* is particularly known for its anti-viral capabilities, including those associated with arthropod-vectored human diseases (Hedges et al. 2008, Moreira et al. 2009, Fenton et al. 2011, Hoffmann et al. 2011, Chrostek et al. 2013), investigating whether *wPni* impacts the transmission dynamics of BBTV may have important implications for pest management strategies and sustainable food production.

## Acknowledgments

We thank Dr. John Thomas (Department of Agriculture, Fisheries, and Forestry (Australia) and Dr Ross Miller (Western Tropical Research Center; University of Guam) for providing aphids for this study. We thank Dr. Susan Halbert for her help in finding *Pentalonia* aphids in Gainesville, FL. We also thank April Greenwell, Shizu Watanabe and Jessika Santamaria for their help with collecting and processing the aphid samples and data curation. The research was funded in part by USDA NIFA titled: Investigating the Phenotypic Effect of *Wolbachia* in Parthenogenetic Aphid Populations, NSF award #1754302 to KMO, and by the NSF Postdoctoral Research Fellowships in Biology Program Grant #2109582 awarded to CHVH.

